# Solving the Diagnostic Odyssey with Synthetic Phenotype Data

**DOI:** 10.64898/2026.03.19.712946

**Authors:** Gianlucca Colangelo, Marcelo Martí

## Abstract

The space of possible phenotype profiles over the Human Phenotype Ontology (HPO) is combinatorially vast, whereas the space of candidate disease genes is far smaller. Phenotype-driven diagnosis is therefore highly non-bijective: many distinct symptom profiles can correspond to the same gene, but only a small fraction of the theoretical phenotype space is biologically and clinically plausible. When a structured ontology exists, this constraint can be exploited to generate realistic synthetic cases. We introduce GraPhens, a simulation framework that uses gene-local HPO structure together with two empirically motivated soft priors, over the number of observed phenotypes per case and phenotype specificity, to generate synthetic phenotype–gene pairs that are novel yet clinically plausible. We use these synthetic cases to train GenPhenia, a graph neural network that reasons over patient-specific phenotype subgraphs rather than flat phenotype sets. Despite being trained entirely on synthetic data, GenPhenia generalizes to real, previously unseen clinical cases and outperforms existing phenotype-driven gene-prioritization methods on two real-world datasets. These results show that when patient-level data are scarce but a structured ontology is available, principled simulation can provide effective training data for end-to-end neural diagnosis models.

## 1. Introduction

Rare diseases collectively affect an estimated 263–446 million people worldwide, yet establishing a molecular diagnosis remains difficult even in the era of routine sequencing [15; 3]. In practice, clinicians often begin not with a complete description of disease expressivity, but with a sparse set of positive findings encoded as Human Phenotype Ontology (HPO) terms [13; 14]. The diagnostic task is therefore to map an incomplete phenotype profile to a causal gene among thousands of candidates. This inference problem is difficult because the observed phenotype set is small, heterogeneous in specificity, and drawn from a large ontology, while only a small fraction of the theoretical phenotype space corresponds to clinically plausible presentations.

HPO makes this problem structured but not simple. Individual phenotype terms, especially broad terms high in the ontology, are associated with many genes, so phenotype– gene relationships are fundamentally many-to-many rather than one-to-one (Supplementary Fig. S1). Existing methods such as Phen2Gene and LIRICAL leverage ontology structure and phenotype specificity to score term-level matches or phenotype-set similarity [24; 17; 20]. These approaches are effective, but they generally aggregate evidence across observed findings rather than explicitly modeling interactions among phenotypes within a patient. This limitation is important because gene-associated phenotypes often span multiple top-level HPO categories rather than remaining confined to a single branch of the ontology; in our study universe, this is true for 93.3% of genes with at least one mapped phenotype category (Supplementary Fig. S2). Rare-disease diagnosis therefore calls for representations that capture not only phenotype specificity, but also the joint structure of co-occurring phenotypes within the ontology.

The central bottleneck is therefore not only gene–phenotype knowledge, but case-level data. For a fixed gene, the phenotypes aggregated across reports define a local phenotype space, whereas an individual clinical record contains only a few of those findings. The number of plausible phenotype profiles is thus combinatorially large even when the number of observed cases is small. Simply sampling arbitrary HPO terms does not solve this problem, because most combinations are biologically implausible and clinically unrealistic. What is needed is a way to expand the training distribution while preserving gene-local ontology structure and the empirical regularities of real clinical records.

We address this problem with GraPhens, a simulation framework that generates synthetic phenotype–gene cases grounded in HPO structure and guided by two empirically motivated soft priors: the number of observed phenotypes per case and phenotype specificity in the ontology. We use these simulated cases to train GenPhenia, a graph neural network that operates on patient-specific phenotype subgraphs rather than flat phenotype sets. Across real clinical benchmarks, GenPhenia trained entirely on synthetic cases outperforms existing phenotype-driven prioritization methods. More broadly, these results suggest that principled simulation can make end-to-end representation learning feasible when real patient-level data are sparse but ontologically structured. Figure 1 provides a visual overview of the clinical motivation, simulation strategy, model, and evaluation setup.

**Figure 1.**
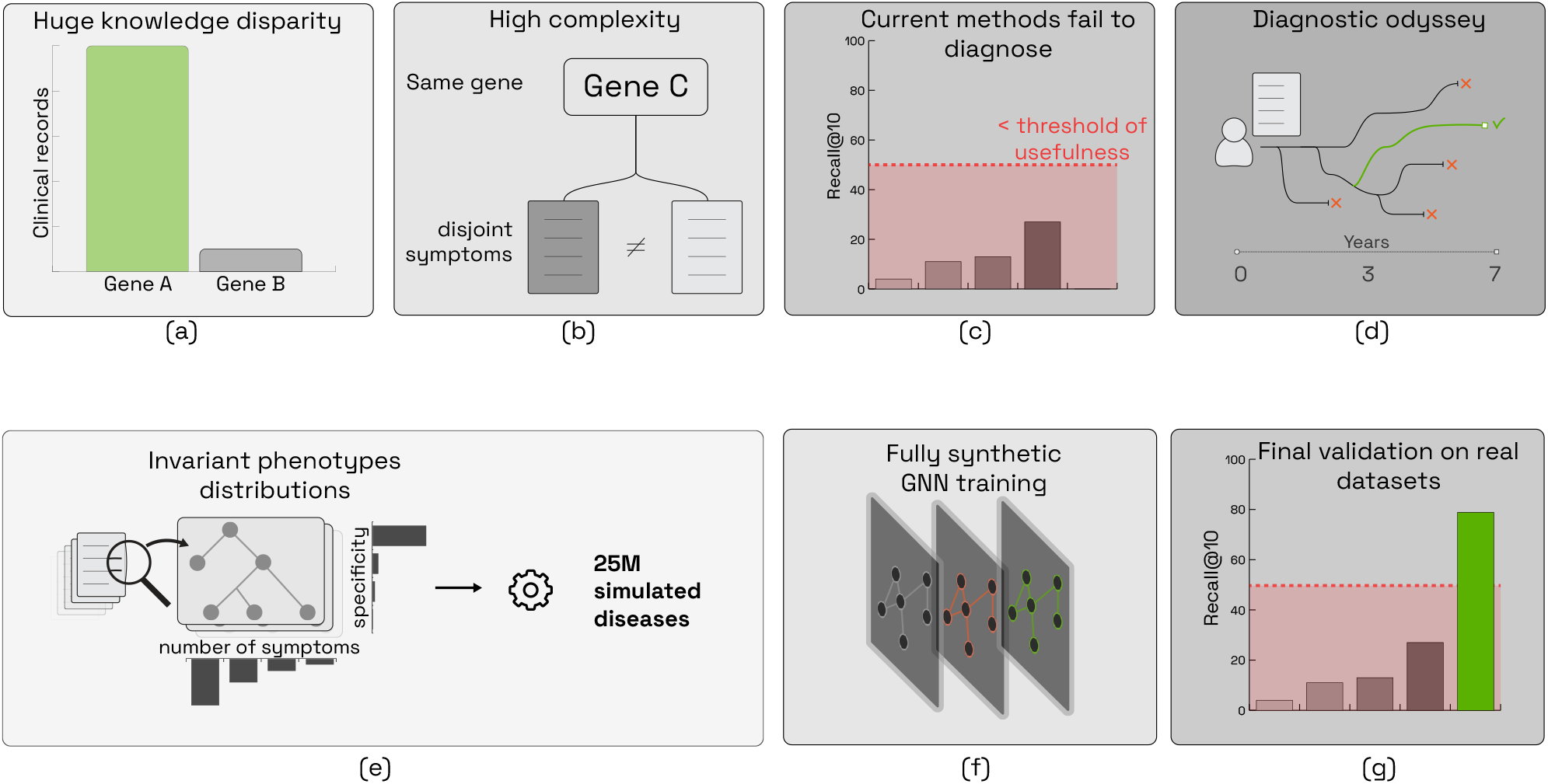
(a) Genetic diseases have a huge disparity in their studies. (b) Also they are highly complex, and same genes can produce completely different set of symptomes, (c) for these reasons, current models fail to predict which genes cause symptoms, all of this ends in a diagnostic odyssey (d). In this work, we present a simulation strategy (e) and train a GNN with fully sinthetic data (f) to finally validate both our simulation strategy and that the model generalizes to real datasets (g).

## 2. Methods

### 2.1. The Human Phenotype Ontology

**The Human Phenotype Ontology (HPO)** provides a standardized vocabulary for describing phenotypic abnormalities observed in human disease and has become a central resource for phenotype-driven rare-disease analysis [13; 14]. HPO is organized as a directed acyclic graph (DAG) in which nodes represent phenotype terms and edges encode “is-a” relations between more general and more specific abnormalities. This structure allows clinical observations to be represented at different levels of granularity, from broad categories close to the root to highly specific findings deeper in the ontology.

Fig. 2 highlights the ontology properties that are most relevant for our framework. First, the HPO release used in this study contains a large space of phenotype terms and gene-linked annotations, providing broad coverage of raredisease presentations. Second, phenotype annotations for a gene are not spread uniformly across the ontology. Instead, for each gene *g*, the associated phenotype set 𝒫_*g*_ occupies a localized region of the graph defined by related terms and their hierarchical neighbors.

**Figure 2.**
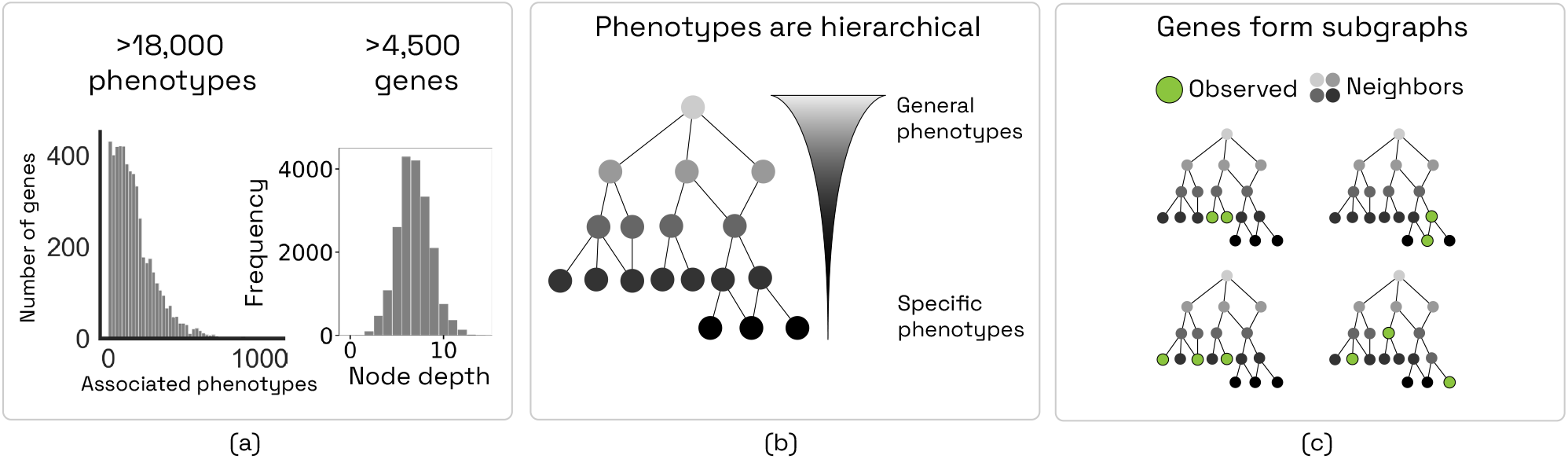
Visual description of the Human Phenotype Ontology. It shows the scale of the ontology and its gene annotations, the distribution of node depths that separates general from specific phenotypes, and the fact that gene-associated phenotypes occupy localized subgraphs rather than the full ontology.

These ontology properties are essential for simulation. A realistic synthetic case should not be produced by sampling arbitrary phenotype terms from the full ontology. Rather, it should be generated from the local phenotype space of a gene while preserving both the number of observed phenotypes and their specificity in the ontology. This ontology-grounded view motivates the simulation strategy introduced in the next subsection.

### 2.2. Simulation strategy

GraPhens simulates phenotype sets by combining a gene-local phenotype space with two empirical priors estimated from rare-disease datasets: the number of observed phenotypes per case and phenotype specificity in HPO (Fig. 3). We denote these priors by *D*_*n*_ and *D*_*s*_, respectively.

**Figure 3.**
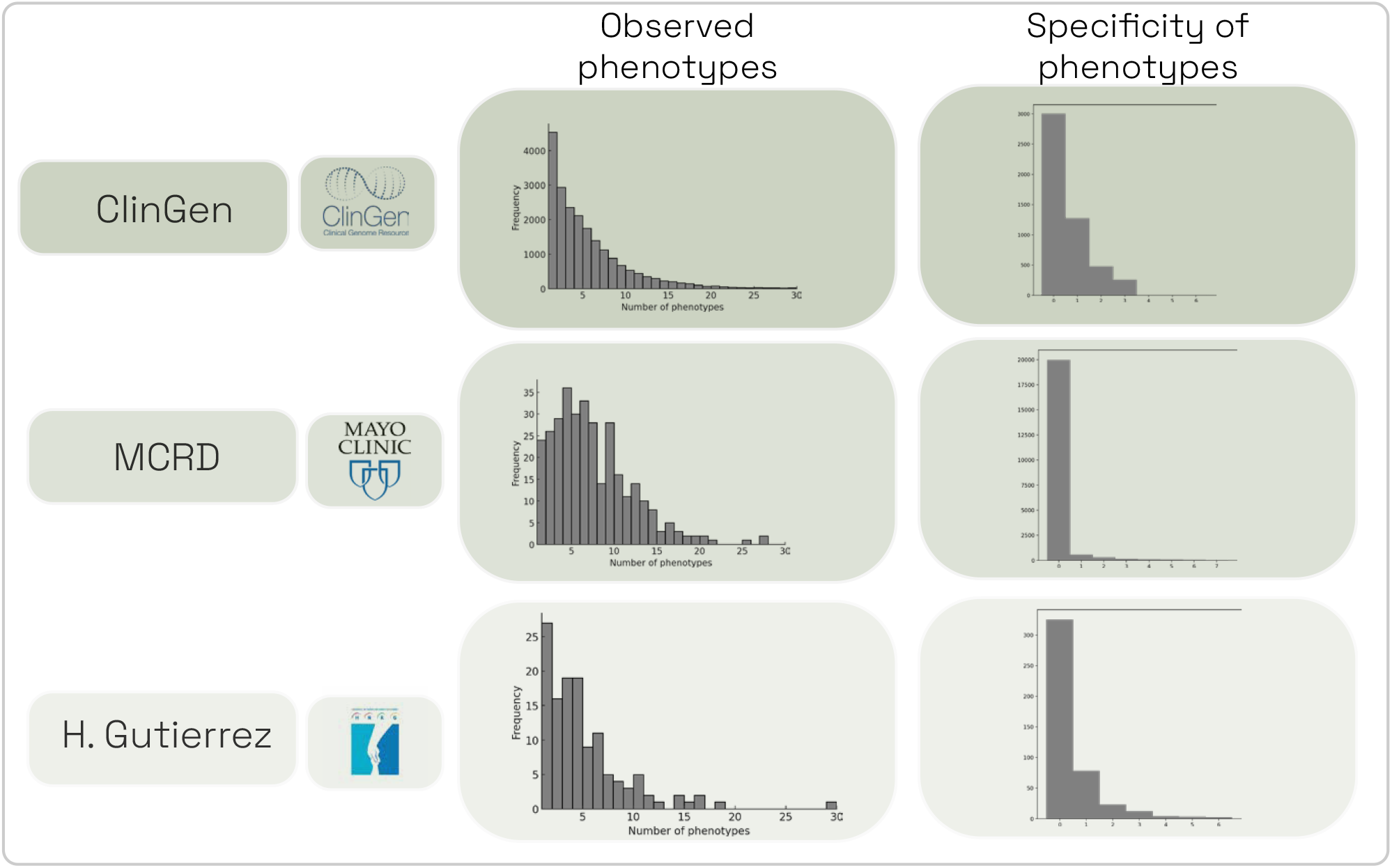
Empirical priors used for simulation. In the rare-disease datasets we examined, the empirical prior over observed phenotype count per case, *D*_*n*_, and the empirical prior over phenotype specificity, *D*_*s*_, exhibit broadly similar shapes. GraPhens uses these empirically motivated soft priors to simulate clinically plausible phenotype sets.

Given a gene *g* with annotated phenotype set 𝒫_*g*_, let 𝒜(*p*) denote the set of ancestors of phenotype *p* in HPO, including the root term `HP:0000001 All phenotypes`. We define the gene-local phenotype space as

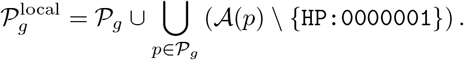

Thus, 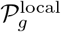 contains the phenotypes directly associated with *g* together with their ontology-based generalizations, but excludes unrelated terms elsewhere in HPO. The universal root term is added deterministically only when constructing the case graph. We first sample the case size *n* ∼ *D*_*n*_. We then draw target specificity values *s*_1_, …, *s*_*n*_ ∼ *D*_*s*_ and, for each target, select a phenotype 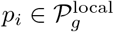 whose ontology depth is compatible with the sampled specificity. The resulting simulated phenotype set is

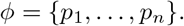

This factorization separates global case statistics from gene-specific phenotype content: *D*_*n*_ and *D*_*s*_ determine how many findings are observed and how specific they are, whereas 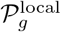 determines which findings can plausibly co-occur for gene *g*. Figure 4 and Algorithm 1 provide complementary summaries of the procedure.

**Figure 4.**
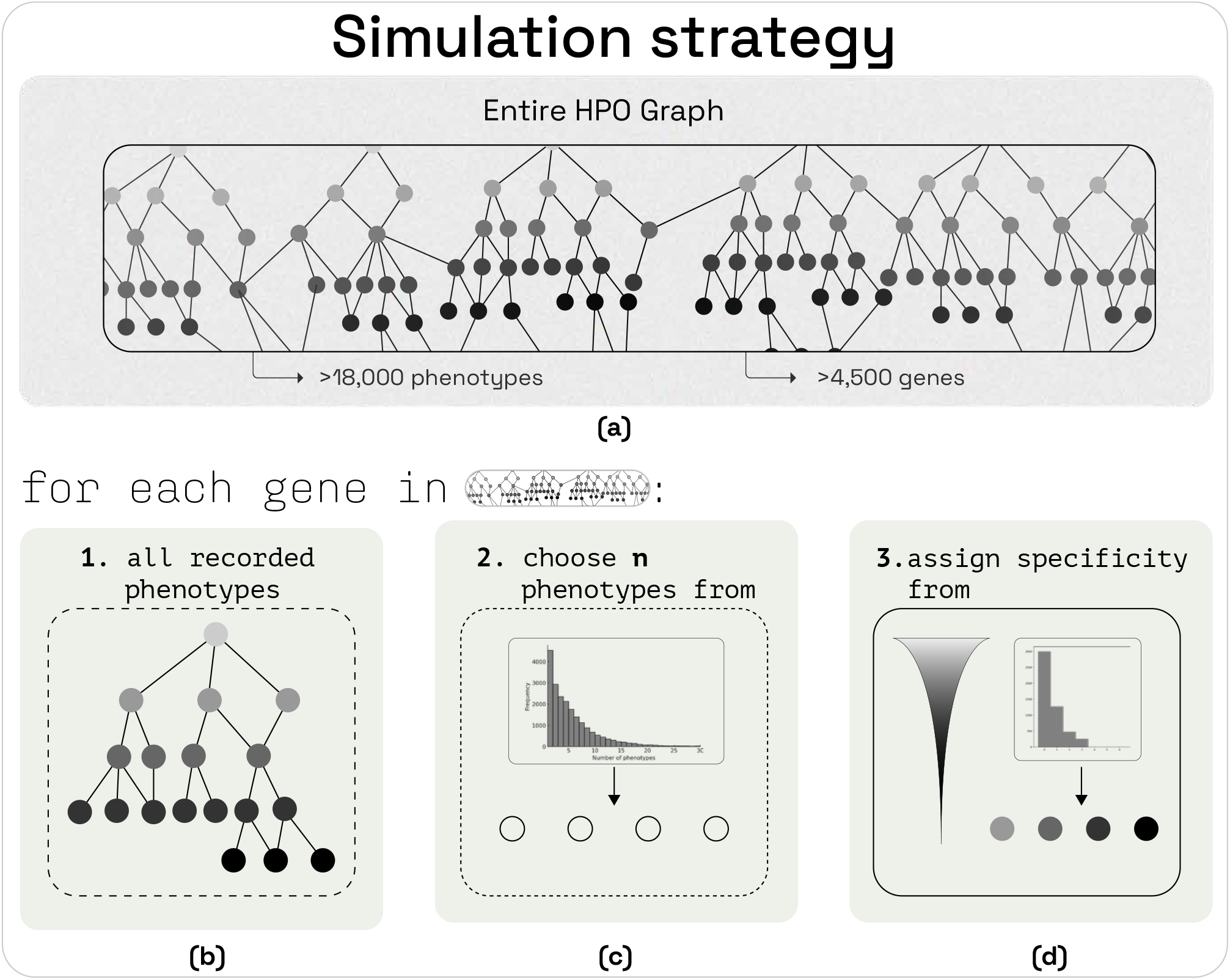
Visual summary of the ontology-grounded simulation procedure. For a selected gene *g*, GraPhens samples the number of observed phenotypes from *D*_*n*_ and samples phenotype-specificity targets from *D*_*s*_. Phenotypes are then drawn from 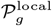, defined as the annotated phenotypes of *g* together with their non-root ontology ancestors; the root term is added later during graph construction. This produces a synthetic phenotype set that is novel, gene-consistent, and constrained by the observed statistical structure of real clinical datasets.

#### Algorithm 1

Ontology-grounded phenotype simulation for gene *g*

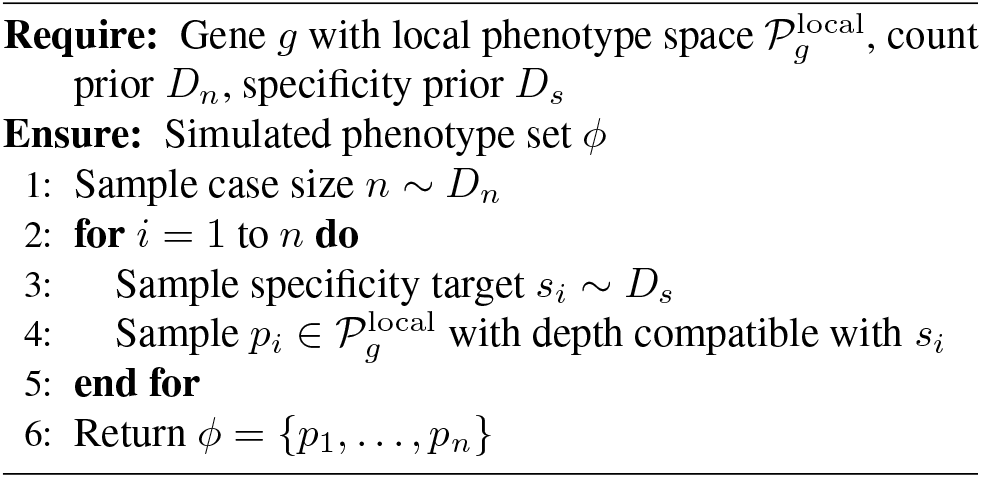

#### 2.2.1. Combinatorial size of the constrained phenotype space

The relevant search space for simulation is not the set of all subsets of HPO, but the set of phenotype profiles that are jointly consistent with gene-local ontology support, the number of observed phenotypes, and phenotype specificity. Let ℒ denote the set of ontology depth bins, and let

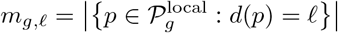

be the number of candidate phenotypes available for gene *g* at depth *ℓ*, where *d*(*p*) is the depth of phenotype *p* in HPO. For a case with *n* observed phenotypes and depth histogram **k** = (*k*_*ℓ*_)_*ℓ*∈ℒ_, satisfying ∑ _*ℓ* ∈ℒ_*k*_*ℓ*_ = *n*, the number of ontology-consistent phenotype profiles is

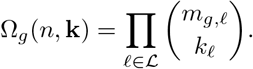

Thus, even after conditioning on gene identity, phenotype count, and phenotype specificity, the space of plausible cases remains combinatorially large. GraPhens therefore defines a constrained forward distribution rather than a finite catalogue of cases. Importantly, this does not make diagnosis trivial: the clinical task is the inverse problem of mapping a sparse phenotype profile *ϕ* back to a causal gene, and multiple genes can occupy overlapping regions of phenotype space through shared general or moderately specific terms. This is precisely why a learned representation that compresses structured phenotype graphs remains necessary even when the generative constraints are known. A worked example and extended discussion are provided in the Supplementary Material.

### 2.3. High-level architecture

#### 2.3.1. Graph augmentation

For each case (simulated or clinical) we generate an HPO-subgraph *G* = (*V, E*) that captures both observed and contextual phenotypes. Starting from the set of positive phenotypes *ϕ*, we form the ancestor closure

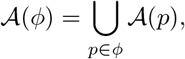

where 𝒜 (*p*) includes the root term `HP:0000001 All phenotypes`. The resulting vertex set

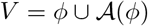

ensures that each case is represented as a rooted ontology subgraph whose depth and branching pattern mirror the native HPO hierarchy.

The native HPO parent–child relation induces a directed edge set *E*_→_ = {(*u, v*) | *u* is parent of *v*}. For message passing, we symmetrize this relation and use the undirected edge set *E* = *E*_↔_ ={{ *u, v*} | (*u, v*) ∈ *E*_→_} . This choice gives the GNN a more flexible receptive field: information can propagate not only along ancestor–descendant chains but also across phenotypes that share a local ancestor, allowing sibling phenotypes to influence one another’s representations through the surrounding ontology structure.

#### 2.3.2. Node features

Each vertex *v* ∈*V* was encoded as the sentence embedding of its HPO definition produced by a biomedical language model. We evaluated three candidate biomedical sentence encoders (see Supplementary Fig. S4) and selected `gsarti/biobert-nli` because, among the candidates, it yielded the least concentrated distribution of pairwise cosine similarities across all 9600 HPO term definitions. We use this statistic only as an intrinsic heuristic for reduced embedding anisotropy and improved discriminability, properties that have been linked to higher-quality sentence representations [5; 21; 6]. Its 768-dimensional embeddings constitute the node-feature matrix *X* ∈ ℝ|*V* |×768. In this way we achieve a richly annotated, variable-sized patient graph that retains ontological structure while supplying continuous clinical features.

### 2.4. Model

The model, GenPhenia, is a graph neural network classifier that receives the variable-sized HPO subgraph of a case and outputs a distribution over 5 229 candidate causal genes (Fig. 5). The architecture consists of three GCN convolutional blocks [11], each comprising a graph convolution, batch normalization, ReLU activation, and dropout (*p*=0.25). The first block projects the 768-dimensional BioBERT node embeddings into 512 hidden channels; the subsequent two blocks preserve this dimensionality. Node-level representations are then aggregated into a single graph-level vector via an attention-gated pooling mechanism that learns to weight phenotype nodes by their diagnostic relevance. A final linear layer maps the resulting 512-dimensional graph embedding to logits over the gene label space.

**Figure 5.**
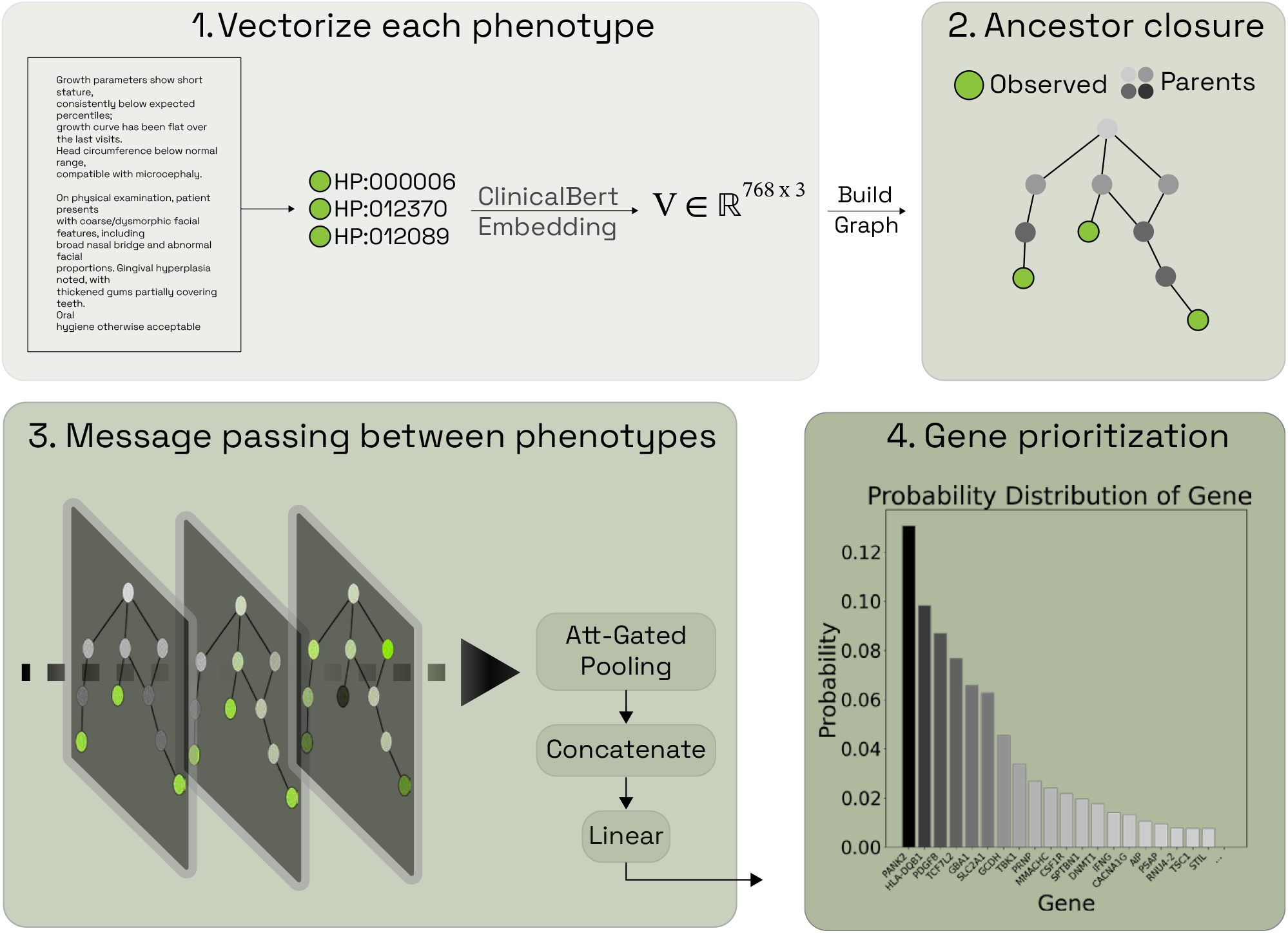
GenPhenia pipeline. Observed HPO terms are encoded with a biomedical language model, assembled into a phenotype subgraph, processed by three GCN blocks (768 →512 →512 →512), aggregated via attention-gated pooling, and classified over 5 229 genes by a linear head.

Because each patient subgraph retains the hierarchical structure of HPO (Section 2.3.1), the three-layer GCN stack gives every node a receptive field that reaches several levels of the ontology. Using the undirected edge set *E*_↔_ also allows information to move between phenotypes that share a local ancestor, so the learned representation of a general term such as “Abnormality of the musculoskeletal system” can be contextualised differently depending on whether the patient’s observed descendants include “Arachnodactyly” or “Brachydactyly.” This lateral exchange among related phenotypes would not occur under a strictly parent-to-child message-passing scheme.

### 2.5. GraPhens: A framework to simulate ontology-grounded diseases

We implemented the simulation strategy described above as GraPhens, an open-source Python library that automates the full pipeline from HPO phenotype identifiers to GNN-ready graph datasets.^1^ The framework manages automatic versioning of the HPO ontology by querying the official release repository for updates, ensuring that all experiments use a consistent term vocabulary. Gene–phenotype annotations are loaded from the HPO Annotation (HPOA) resource, which aggregates phenotype evidence from OMIM, Orphanet, and ClinGen-curated gene–disease validity assertions. Simulation and graph-augmentation strategies are composable modules, so that the empirical-distribution simulation used in our main experiments and the uniform-distribution ablation control described in Section 2.6 share the same phenotype-selection and graph-construction code, differing only in the sampling distributions *D*_*n*_ and *D*_*s*_.

### 2.6. Ablation design

To disentangle the contributions of the simulation strategy from those of the model architecture, we ablate along two axes—the simulation regime used to generate training data and the model family used for classification—and evaluate all four combinations (Table 1).

**Table 1.** 2 ×2 ablation design. Rows vary the model; columns vary the simulation regime.

| | Naive simulation<br>$n \sim \text{Unif}\{1, \dots, 30\}, s \sim \text{Unif}\{0, \dots, 8\}$ | Realistic simulation<br>$n \sim D_n, s \sim D_s$ |
| --- | --- | --- |
| FNN (mean-pool) | FNN+naive | FNN+realistic |
| GNN (graph) | GNN+naive | GNN+realistic |

#### Architecture factor

The GNN described in Section 2.4 is compared against a feedforward neural network (FNN) baseline that receives the same BioBERT embeddings but discards ontology structure. For each case the FNN mean-pools the 768-dimensional embeddings of the observed phenotypes into a single vector and classifies it with a multilayer perceptron. Because the FNN input is a fixed-dimensional summary statistic of the phenotype set, its representation is directly sensitive to shifts in the number of observed phenotypes and their specificity: a mismatch between training and evaluation distributions in these two quantities constitutes a first-order domain shift.

#### Simulation factor

The empirical-distribution simulation introduced in Section 2.2 is compared against a naive baseline in which both the case size *n* and the per-phenotype specificity target *s* are drawn from uniform distributions (*n* ∼Uniform {1, …, 30}, *s* ∼Uniform {0, …, 8}). The naive regime is still gene-anchored—phenotypes are selected from 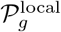 —but it overrepresents extreme case sizes and overly general terms relative to real clinical records. Both simulation regimes share the same phenotype-selection and graph-construction code within GraPhens, differing only in the sampling distributions *D*_*n*_ and *D*_*s*_.

### 2.7. External phenotype-only benchmarks

We compare GenPhenia against four published phenotype-only gene-prioritization methods. PCAN evaluates phenotype consensus in the signaling neighborhood of a candidate gene [8]. Phen2Gene aggregates precomputed HPO-term-specific gene rankings from the HPO2Gene Knowledgebase [24]. CADA performs prioritization on a case-enriched phenotype–gene knowledge graph [16]. PPAR uses embeddings from a clinical knowledge graph linking genes, HPO terms, and Gene Ontology annotations [7].

We assess transfer on two external clinical cohorts. The first is the public DDD cohort, originally described by Wright et al. [22] and later reused in the recent PPAR evaluation [7]. The second is the Mayo Clinic rare-disease (MCRD) cohort reported in the PPAR study [7]. Figure 6 summarizes Recall@N on these two cohorts.

**Figure 6.**
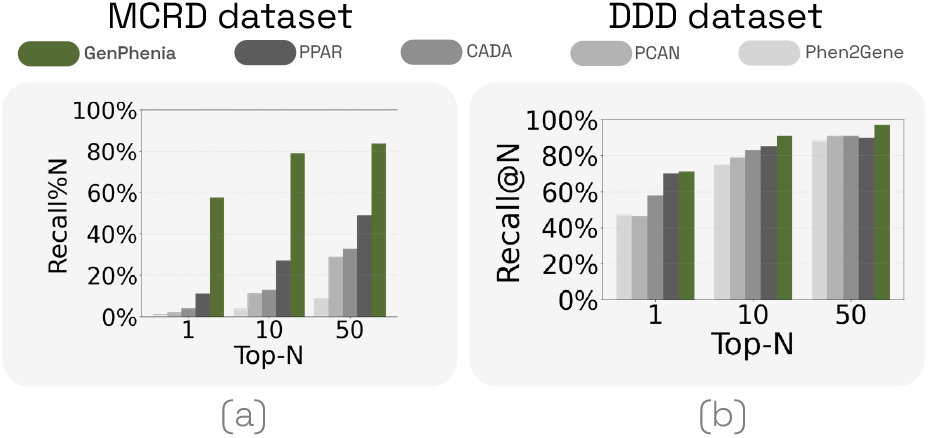
Comparison with four published phenotype-only gene-prioritization methods on the DDD and MCRD cohorts. Bars report Recall@N for GenPhenia, PPAR, CADA, PCAN, and Phen2Gene.

## 3. Results

### 3.1. GenPhenia trained on synthetic cases transfers to real clinical cohorts

Figure 6 reports performance on these two external cohorts. Despite being trained entirely on synthetic cases generated by GraPhens, GenPhenia achieves the highest Recall@N on both benchmarks. On the public DDD cohort, GenPhenia reaches 91% Recall@10, compared with 85% for PPAR, 83% for PCAN, 79% for Phen2Gene, and 75% for CADA. On MCRD, GenPhenia reaches 78.9% Recall@10, compared with 27% for PPAR, 13% for CADA, 11% for PCAN, and 4% for Phen2Gene. It also maintains the strongest ranking performance across the other reported cutoffs shown in Fig. 6. These results indicate that ontology-grounded synthetic training is sufficient to learn phenotype–gene structure that transfers to external clinical cases.

In the main training regime, we generated 5,000 synthetic cases per gene, and none of the simulated phenotype sets exactly matched a real evaluation case. This rules out record-level memorization as an explanation for the benchmark performance and is consistent with the combinatorial analysis in Section 2.2.1: even when a real case lies within the genelocal phenotype support 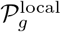, the number of ontology-consistent phenotype subsets remains large enough that exact reproduction is unlikely.

### 3.2. Empirical phenotype priors matter primarily for the non-graph baseline

Figure 7 summarizes the 2 ×2 ablation using training sets with 2,500 simulated cases per gene. Replacing the FNN with the GNN yields the largest improvement in Recall@1 under both simulation regimes. The effect of the simulation strategy is strongest for the FNN: realistic simulation increases Recall@1 from approximately 0.06 to 0.27, whereas the corresponding change for the GNN is small, from approximately 0.42 to 0.43. Thus, matching the empirical priors *D*_*n*_ and *D*_*s*_ is important for the FNN baseline, while the GNN remains comparatively stable even when trained on the naive simulation regime.

**Figure 7.**
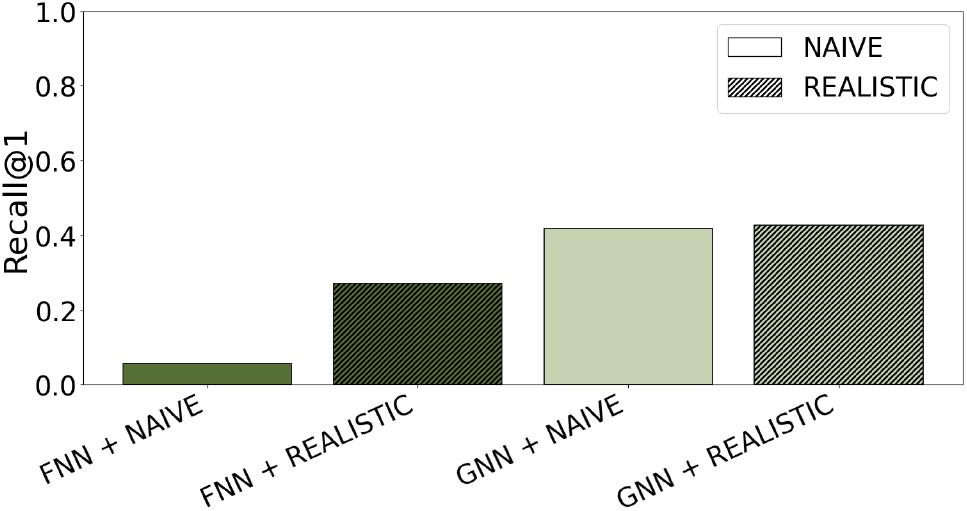
Ablation results for the 2*×* 2 study design. Bars report Recall@1 for the four combinations of architecture and simulation regime: FNN+naive, FNN+realistic, GNN+naive, and GNN+realistic.

A stratified analysis by phenotype count and mean HPO depth is provided in Supplementary Fig. S3. Consistent with prior work, performance improves as phenotype sets become larger and more specific [12; 20].

## 4. Discussion

The central contribution of GraPhens is to exploit the structure of HPO to generate phenotype–gene cases that remain clinically plausible. When a structured ontology exists, this constraint can be used to generate realistic synthetic cases: in our setting, gene-local HPO neighborhoods delimit which phenotypes can co-occur, while the empirical priors *D*_*n*_ and *D*_*s*_ regularize how many findings are observed and how specific they are. The resulting training distribution is therefore novel at the record level while remaining close to the clinically relevant region of phenotype space. GenPhenia’s ability to transfer from these simulated cases to previously unseen real patients (Fig. 6) suggests that the simulator preserves task-relevant phenotype–gene structure rather than record-level identity [9; 23; 4].

The 2×2 ablation also clarifies the relation between empirical priors and architectural inductive bias. Realistic simulation substantially improves the FNN baseline, but has only a minimal effect on the GNN. This is expected because the FNN reduces each case to a mean-pooled embedding, so shifts in phenotype count and specificity directly perturb its input distribution. By contrast, the GNN remains strong even under naive simulation, which suggests that it is comparatively robust to misspecification of the empirical priors *D*_*n*_ and *D*_*s*_. Importantly, the naive regime is not prior-free: phenotypes are still sampled from the gene-local ontology support 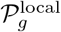 . The ablation therefore indicates that accurate matching of the empirical phenotype-count and specificity marginals is far less critical for the GNN once it can exploit phenotype structure through message passing on the HPO graph.

This interpretation is consistent with prior work on relational inductive biases and combinatorial generalization in graph networks. Graph-based models have been shown to transfer across changes in the number of entities and relations in physical systems, to learn useful representations for previously unseen nodes and graphs, and to generalize across both graph size and input distribution in structured reasoning tasks [1; 2; 18; 10; 19]. Our setting is different from those benchmarks, but it shares the same core ingredient: explicit relational structure. Here, HPO provides that structure, GraPhens uses it to sample realistic synthetic cases, and GenPhenia exploits patient-specific phenotype subgraphs through multi-hop message passing over the ontology. In that sense, the results support the broader view that explicit structure and flexible learning can improve generalization when training data are limited but the domain is richly organized.

One open question remains. We do not yet know whether the cross-dataset similarity of phenotype count and specificity reflects biological regularities, clinical ascertainment, or both. Resolving that question would sharpen the theoretical scope of GraPhens, even though the observed similarity of these marginals is already sufficient to make the simulator useful in clinical practice.

## Supplementary Material

This section contains four supplementary components referenced in the main text: an extended discussion of the combinatorial size and inverse ambiguity of the constrained phenotype space, an intrinsic encoder-selection diagnostic referenced in the node-feature description above, and two ontology-level analyses supporting the Introduction. The ontology-level analyses show how gene association counts vary with phenotype specificity and quantify the breadth of gene-associated phenotypes across top-level HPO categories.

### Combinatorial size and inverse ambiguity of the constrained phenotype space

The search space relevant to simulation is not the unconstrained power set of HPO, but the subset of phenotype profiles that are compatible with a gene’s local ontology neighborhood and with the empirical constraints imposed by real clinical records. Prior work has shown that sparse but highly specific HPO terms carry disproportionate diagnostic signal, whereas randomly selected, less-specific terms degrade gene prioritization performance [20; 24]. Likewise, LIRICAL has shown that parent or grandparent terms provide a realistic model of phenotypic imprecision, which confirms that ontology depth is a meaningful axis of information loss in phenotype-driven diagnosis [17]. These observations imply that clinically plausible cases are jointly constrained by gene-local support, the number of observed findings, and their specificity in HPO.

Let ℒ denote the set of ontology depth bins, and let

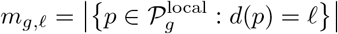

be the number of candidate phenotypes available for gene *g* at depth *ℓ*, where *d*(*p*) is the depth of phenotype *p* in HPO. Because each phenotype can appear at most once in a clinical record, selection within each depth stratum is without replacement. For a case with *n* observed phenotypes and depth histogram **k** = (*k*_*ℓ*_)_*ℓ*∈ℒ_, satisfying ∑ _*ℓ* ∈ℒ_*k*_*ℓ*_ = *n*, the number of ontology-consistent phenotype profiles is

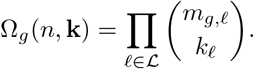

As an illustrative example, suppose that a gene has 25 candidate phenotypes at one depth bin, 40 at a second, and 18 at a third, and that a simulated case draws the depth histogram **k** = (2, 2, 1). Then

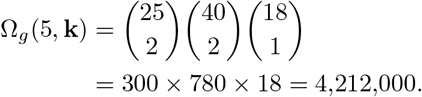

This is a conservative lower bound because it counts only phenotype identity combinations within each specificity stratum and does not incorporate phenotype frequency, age of onset, or higher-order phenotype dependencies. Even so, a single three-bin configuration already yields millions of distinct phenotype profiles for one gene, and thousands of simulated cases per gene explore only a small fraction of this reachable space.

Crucially, knowing the forward constraints that define this space does not solve the diagnostic problem. Diagnosis is an inverse inference problem: given a sparse phenotype profile *ϕ*, the goal is to infer which gene among thousands most plausibly generated it. Multiple genes can induce intersecting feasible phenotype regions, particularly through shared broad or moderately specific phenotypes, so the mapping from phenotype profile to gene remains highly non-bijective. This is the setting in which learned representations are useful: they do not replace the ontology-grounded rules used for simulation, but compress a vast and overlapping structured space into features that improve discrimination among candidate genes.

### Phenotype specificity and propagated gene association counts

**Figure S1.**
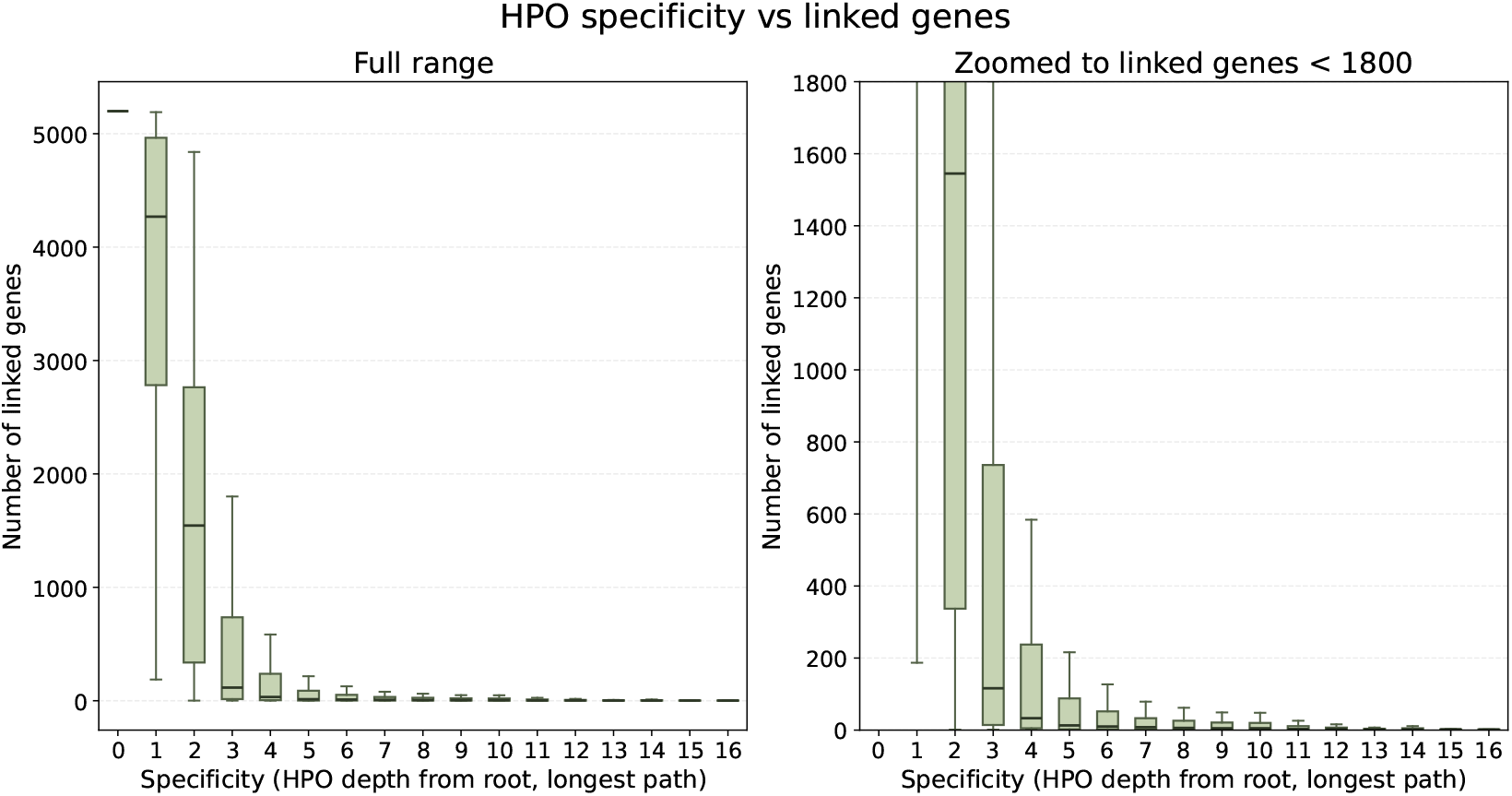
Distribution of propagated gene counts per HPO term stratified by ontology specificity, operationalized as node depth. Each box summarizes how many genes are associated with terms at a given depth after propagating annotations to ancestor terms, which makes the many-to-many phenotype–gene structure explicit: broad, shallow terms are linked to far more genes than specific descendant terms. For example, `HP:0033127 Abnormality of the musculoskeletal system` appears at depth 2 with 3,870 propagated gene associations, whereas `HP:0002661 Painless fractures due to injury` has 8 associated genes and appears at specificity 7.

### Top-level HPO category breadth per gene

To complement the depth-stratified analysis above, we also quantified how broadly each gene’s phenotype annotations spread across the ontology. For every gene–HPO annotation in the study universe, we mapped the annotated term upward to the immediate child category of `HP:0000118 Phenotypic abnormality` and counted the number of unique top-level categories represented per gene. This produces a gene-level multisystem breadth proxy rather than a disease-level clinical definition of syndromic presentation, but it directly captures whether phenotype evidence for a gene remains confined to a single top-level branch or spans multiple branches of HPO.

**Figure S2.**
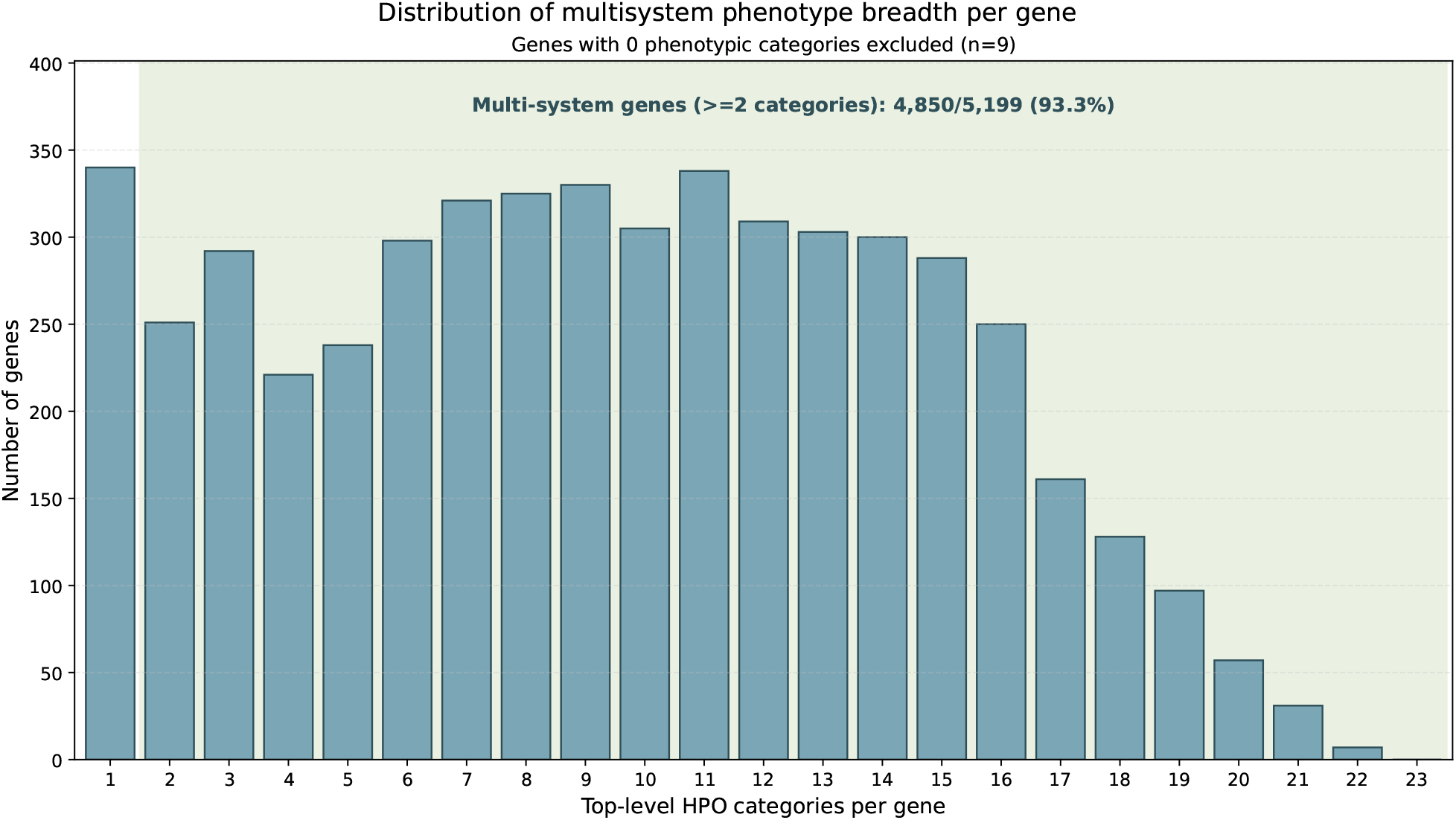
Distribution of top-level HPO category breadth per gene. Each bar reports how many genes are annotated to exactly *k* unique top-level categories, where categories are defined as the immediate children of `HP:0000118 Phenotypic abnormality`. Genes with zero mapped top-level categories (*n* = 9) are excluded. Under this operational definition, a gene is considered multisystem if its annotations span at least two top-level categories; 4,850 of 5,199 genes with at least one mapped category (93.3%) satisfy this criterion. This figure is intended as an ontology-level breadth diagnostic motivating models that capture co-occurring phenotypes across multiple HPO branches. It should not be interpreted as a formal clinical definition of syndromic disease, because these top-level categories include cross-cutting classes such as growth abnormality, constitutional symptom, neoplasm, and prenatal development in addition to organ-system-specific branches.

### Performance by phenotype number and specificity

**Figure S3.**
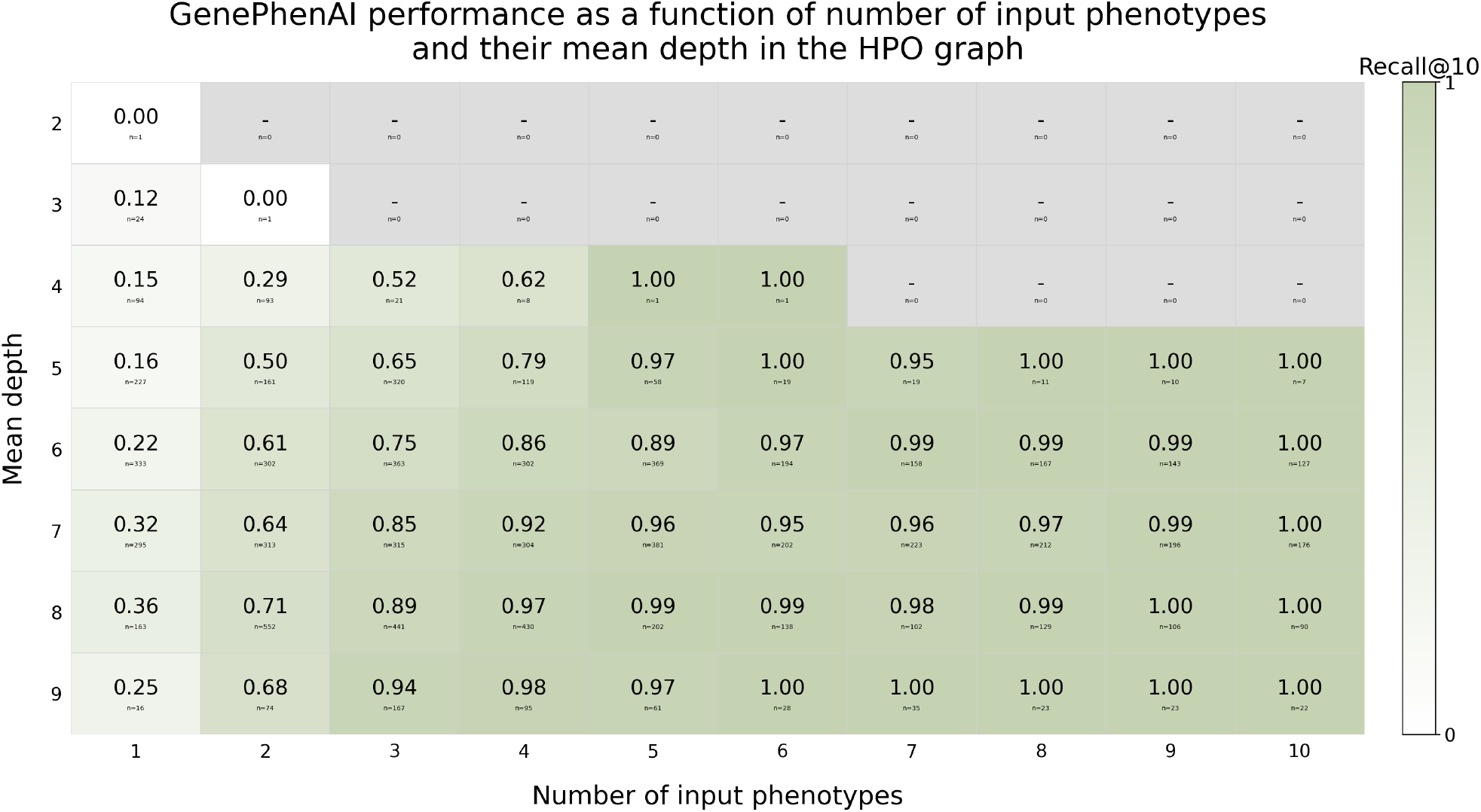
Recall@10 of GenPhenia by number of input phenotypes and mean HPO depth on phenotype subsets derived from 500 ClinVar cases. Each cell shows the mean Recall@10, and smaller text reports the number of evaluated subsets in that bin; cells marked “-” contain no subsets. Higher performance concentrates in bins with larger and more specific phenotype sets.

Figure S3 stratifies Recall@10 by the number of input phenotypes and their mean depth in HPO using phenotype subsets derived from a random subsample of 500 ClinVar cases. Performance is weakest when the phenotype set is both small and shallow in the ontology: subsets with one or two phenotypes and mean depth 2–3 yield very low recall or are absent. Recall@10 then increases steadily as the phenotype sets become larger and more specific. Once cases contain at least five phenotypes with mean depth at least 5, performance is consistently high, and many bins approach or reach 1.00. This trend is consistent with prior work showing that HPO-based prioritization benefits from more specific phenotype information and degrades under phenotypic imprecision [12; 20].

### Intrinsic encoder-selection diagnostic

**Figure S4.**
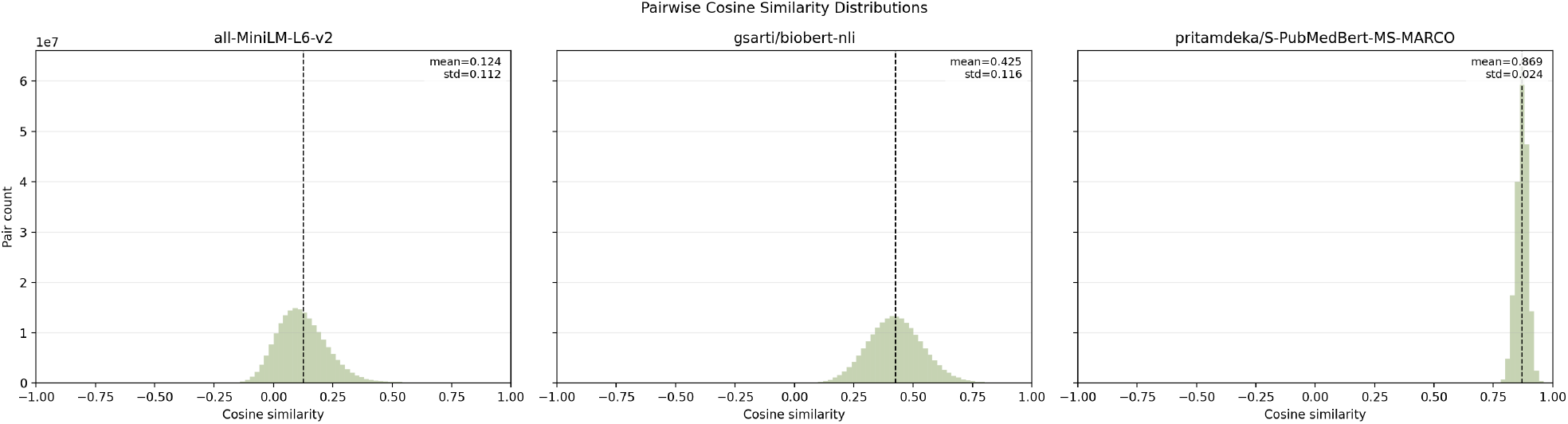
Pairwise cosine-similarity histograms across all 9600 HPO term definitions for the three candidate biomedical sentence encoders. Among the evaluated models, `gsarti/biobert-nli` produced the least concentrated similarity distribution and was therefore selected for the node features used in the main experiments.

## Footnotes

1 Code available at https://github.com/gianluccacolangelo/GraPhens. Public demo available at https://genphenia.xyz/.

## Notes

https://github.com/gianluccacolangelo/GraPhens

https://genphenia.xyz/

### Competing Interest Statement

The authors have declared no competing interest.

